# Downstream promoter interactions of TFIID TAFs facilitate transcription reinitiation

**DOI:** 10.1101/184317

**Authors:** Yoo Jin Joo, Scott B. Ficarro, Luis M. Soares, Yujin Chun, Jarrod A. Marto, Stephen Buratowski

## Abstract

TFIID binds promoter DNA to recruit RNA polymerase II and other basal factors for transcription. Although the TATA-Binding Protein (TBP) subunit of TFIID is necessary and sufficient for in vitro transcription, the TBP-Associated Factor (TAF) subunits recognize downstream promoter elements, act as co-activators, and interact with nucleosomes. Here we show that transcription induces stable TAF binding to downstream promoter DNA, independent of upstream contacts, TBP, or other basal transcription factors. This transcription-dependent TAF complex promotes subsequent activator-independent transcription, and promoter response to TAF mutations in vivo correlates with the level of downstream, rather than overall, Taf1 crosslinking. We propose a new model in which TAFs function as reinitiation factors, accounting for the differential responses of promoters to various transcription factor mutations.

## Introduction

Transcription by RNA polymerase II (RNApII) requires a set of basal initiation factors that recognize core promoter sequences, position the polymerase appropriately, catalyze promoter melting, and mediate the transition from preinitiation (PIC) to stable elongation (EC) complexes (Thomas and Chiang, 2006). Building on decades of biochemistry and molecular genetics, recent structural studies are providing insight into how these factors fit and function together (Hahn and Buratowski, 2016). However, some aspects of initiation factor function remain unclear. In particular, the basal factor TFIID has been the subject of much study and debate.

Early template commitment and DNAse I footprinting experiments identified TFIID as the factor that first binds promoter DNA. It contacts the TATA box, as well as DNA downstream near the transcription start site (TSS) and beyond (Nakajima et al., 1988). A single small protein designated TATA-Binding Protein (TBP) is necessary and sufficient for PIC assembly and TFIID transcription activity in vitro (Buratowski et al., 1989; 1988). However, some fraction of TBP in cells is associated with roughly a dozen additional proteins called TBP-Associated Factors, or TAFs for short (reviewed in Thomas and Chiang, 2006). While not required for in vitro transcription, TAFs are generally essential for cell and organism viability. Multiple activities have been proposed for TAFs.

One TAF function is to contact promoter DNA. Very short elements around the TSS and 20-30 bp downstream can contribute to promoter strength (Kadonaga, 2012). These may affect RNApII, TFIIH, or TFIID binding, but the effect of these elements is strongest when there is a weak non-consensus TATA box. The TFIID DNAse I footprint on some promoters extends much further downstream than TBP alone, and several TAFs crosslink at the TSS and beyond (Auty et al., 2004; Burke and Kadonaga, 1997; Horikoshi et al., 1988a; 1988b; Lee et al., 2005; Sypes and Gilmour, 1994). A recent cryo-EM structure shows Taf1, Taf2, and Taf7 directly interacting with the TSS and +20-+30 regions (Louder et al., 2016). Therefore, a subset of TAFs likely mediates extended promoter recognition.

TAFs have also been implicated in transcription activation. In vitro transcription with TBP supports basal transcription, but response to activators is often potentiated by TFIID. At some promoters, DNAse I protection by TFIID downstream of the TATA box is only seen in the presence of an activator, suggesting TFIID is not only a target for activators, but may undergo activator-induced conformation changes (Chi and Carey, 1996; Horikoshi et al., 1988a; 1988b). The TAF "co-activator" model became controversial when early experiments in yeast showed some genes continued expressing after TAFs were inactivated (summarized in Hahn, 1998). In addition, compelling genetic and biochemical data showed many activators function by contacting and recruiting other factors. Activator targets include histone acetyltransferases (HATs) and chromatin remodelers that make promoter DNA more accessible, as well as the Mediator-RNApII complex needed for PIC assembly. Nevertheless, both in vitro and in vivo experiments argue that some activators function via direct contacts with specific TAF subunits (Albright and Tjian, 2000; Chen et al., 2013; Mencía et al., 2002; Papai et al., 2010).

Over time, extensive analyses produced a consensus that at least 90% of yeast genes show reduced expression upon TAF inactivation (Huisinga and Pugh, 2004; Lee et al., 2000). The affected promoters tend to have weaker, non-consensus TATA elements and are more likely to be constitutively expressed. The 10% of promoters that are relatively resistant to TAF mutations are instead strongly affected by mutations in another coactivator, the histone H3 acetyltransferase SAGA. These genes generally have a strong consensus TATA and are more likely to be highly induced under specific conditions. While often over-simplified to categorize each gene as dependent on only one or the other complex, the data actually show combinatorial effects of double mutants at nearly all promoters (Bonnet et al., 2014; Grünberg et al., 2016; Huisinga and Pugh, 2004). To reflect this subtlety, promoters are often referred to as TFIID-dominant or SAGA-dominant, although this categorization doesn’t fully reflect the continuum of promoter responses. Importantly, many activators can function at both promoter classes, arguing that it not simply the specific activator that determines response to TAFs (de Jonge et al., 2016).

TAFs are also likely to mediate communication between TFIID and nucleosomes. Several TAFs contain histone-fold domains, although it remains unclear if they form a structure resembling a nucleosome within TFIID or interact with actual histone proteins (Bieniossek et al., 2013; Selleck et al., 2001). The C-terminal half of Taf1 in higher eukaryotes has two bromodomains and an extended C-terminal domain, i.e. a BET module. In S. cerevisiae, the BET module is encoded by the *BDF1* gene rather than being fused to Taf1 (Matangkasombut et al., 2000). The Bdf1 bromodomains bind specifically to acetylated histone H4 tails (Ladurner et al., 2003; Matangkasombut and Buratowski, 2003). It remains unclear if and how this interaction affects transcription.

Current models for TAF function are premised on TFIID acting before transcription initiation. However, while carrying out a proteomic analysis of RNApII ECs, we unexpectedly discovered a striking transcription-dependent recruitment of TAFs, but not TBP or other basal factors, to DNA well downstream of the TATA box. In vitro transcription reactions suggest this post-initiation TAF complex promotes additional rounds of transcription independently of an upstream activator. The in vivo relevance of these downstream interactions is supported by ChIP-exo data showing a downstream-extended TFIID footprint specifically on TFIID-dominant genes. Based on these data, we propose a new function for the TAF complex as an activator-independent reinitiation factor. This model explains some earlier confusing data about TFIID function and provides a simple explanation in which the level of TFIID-dominance is determined by how efficiently a promoter supports TAF-dependent reinitiation.

## RESULTS

### Transcription-dependent recruitment of TAFs to downstream DNA

We and others have used quantitative mass spectrometry to analyze RNApII initiation complexes assembled in vitro on bead-immobilized templates (Ranish et al., 2004; Sikorski et al., 2012). We extended this approach to elongation complexes (Joo et al., in preparation). DNA templates containing five Gal4 binding sites and the *CYC1* core promoter upstream of a G-less cassette were immobilized on beads and incubated with nuclear extract. In the presence of ATP, UTP, CTP, and the chain terminator 3’-O-me-GTP, ECs extend to the end of the G-less cassette and stall. Digestion with restriction enzyme SspI cuts 30 bp downstream of the TATA box to release ECs bound to downstream DNA, while any remaining PIC components stay associated with upstream DNA on the bead (Fig 1A).

**Figure 1.**
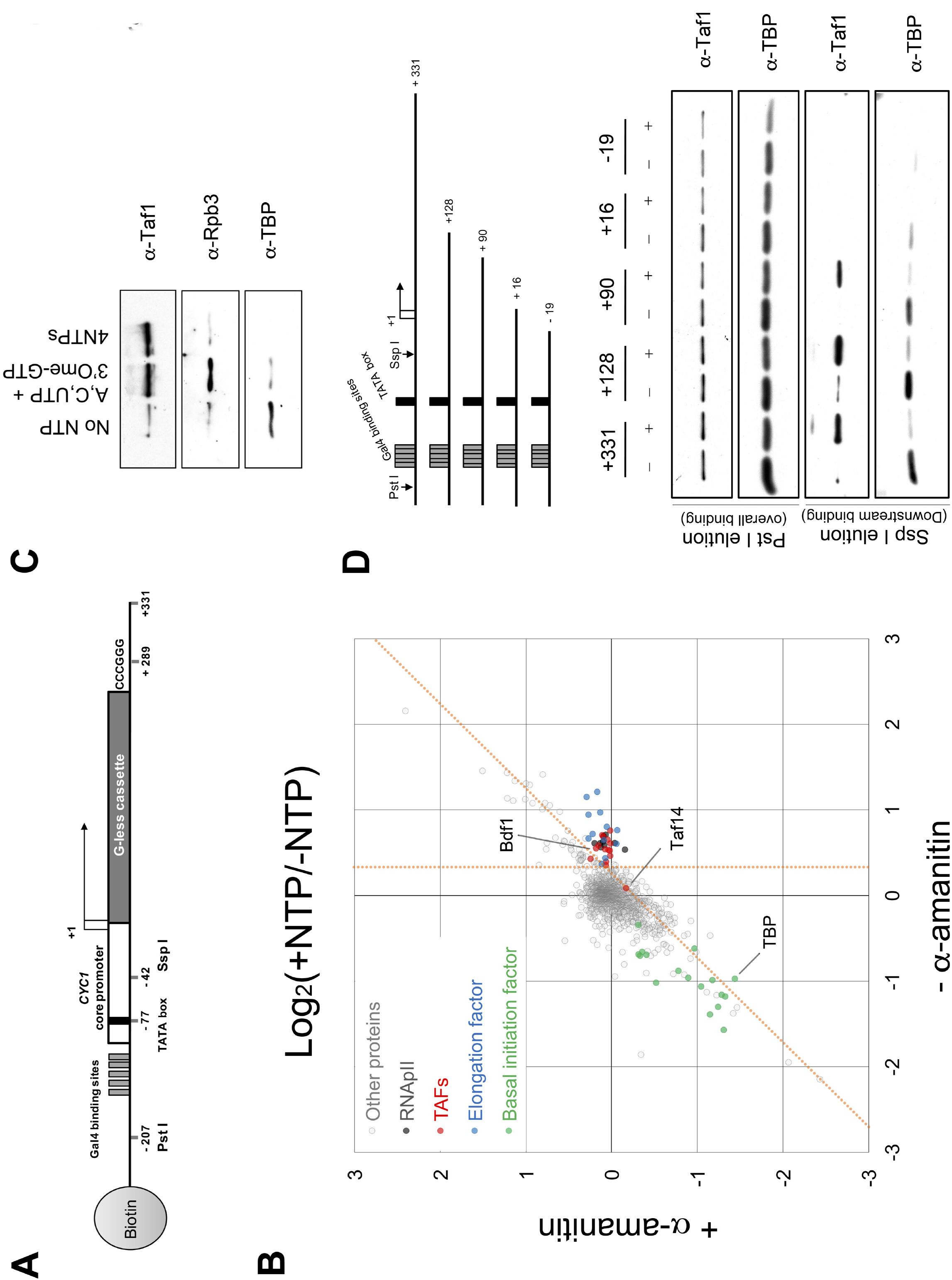
Transcription-mediated TAF association with downstream DNA. A. Schematic of transcription template used to isolate RNApII ECs as described in Methods. Downstream bound proteins are eluted using Ssp I endonuclease, which cuts 35 nts downstream of the TATA box but upstream of the first TSS. Total proteins are eluted with Pst I (see **Fig S1D**). B. Scatter plot comparing NTP-dependent protein binding to downstream DNA (log2 of + NTP /- NTP ratio) in the presence (y-axis) or absence (x-axis) of α-amanitin. Each circle represents one protein (1511 total), with the value calculated from the sum of all peptide signals identified for that protein. Proteins in specific subgroups are color coded as noted. Proteins specifically dependent on transcription are defined as those to the right of the vertical orange line (95% confidence level for enrichment with NTPs) on the x-axis and below the diagonal orange line (95% confidence level cutoff for reduction of NTP enrichment by alpha-amanitin). See also **Fig S1A-C**, and **Table S1**. C. Immunoblot of proteins eluted from immobilized templates with Ssp I before treatment with NTPs, after 15 minutes incubation with A, C, and UTP plus 3’O-meGTP (to stall elongation complexes), or with all four NTPs to allow RNApII runoff. D. TBP and Taf1 binding on immobilized templates of different lengths. Upper panel shows schematic of template series analyzed as in part B. The middle panels show immunoblots of total bound proteins eluted with Pst I, while the lower panels show downstream binding proteins eluted with Ssp I.

Quantitative mass spectrometry revealed multiple proteins associated with downstream DNA specifically in the presence of NTPs. To distinguish which were dependent upon transcription rather than simply NTP hydrolysis, a parallel reaction with the elongation inhibitor α-amanitin was analyzed. Statistically significant enrichment (Fig 1B, **S1A-C, and Table S1**) of RNApII subunits (black circles) and known elongation factors such as PAF complex and Spt5 was observed (blue circles), showing that bona fide ECs were isolated. A detailed description of these proteomic results will appear elsewhere (Joo et al., in preparation).

We were surprised to also detect very clear transcription-dependent enrichment of all known TAF proteins (red circles) including Bdf1, on the released downstream DNA. The one exception was Taf14/Tfg3, a YEATS domain protein also found in multiple yeast complexes, but not mammalian or Drosophila TFIID (Thomas and Chiang, 2006). Enrichment of all Tafs except Taf14 and Taf10 (a small protein in both TFIID and SAGA) was blocked by alpha-amanitin (Fig 1B, **S1C**). Surprisingly, TBP behavior differed from TAFs. The already low signal for downstream binding of TBP and other basal transcription factors (likely PICs formed at weak cryptic TATAs in the G-less cassette) was further reduced by NTPs, even when transcription was inhibited (green circles). The increase in TAFs is less pronounced when the entire transcription template is released by Pst I digestion, arguing that downstream TAFs may be transferred from upstream (Fig 1D, **S1D**). Similar downstream TAF enrichment is seen on either naked or chromatinized templates (**Fig S1E**). From these data, we conclude that release of RNApII from the PIC into elongation leaves TAF proteins stably associated with downstream promoter DNA, independent of TBP remaining bound to the TATA box.

Many results argue TAFs do not associate with ECs. First, crosslinking experiments detect TAFs near promoters but not downstream (Rhee and Pugh, 2012). Second, the curated literature (www.yeastgenome.org) contains no reports of TAF mutants exhibiting 6-aza-uracil or mycophenolic acid sensitivity, phenotypes often associated with elongation defects. Finally, affinity purifications of TAFs and elongation factors failed to observe physical association (Gavin et al., 2006; Krogan et al., 2006). To verify that TAFs are not associated with ECs in our system, we compared a reaction with U, C, and ATP plus 3’-O-me-GTP, where ECs stall at the end of the G-less cassette, to a reaction with all four NTPs, allowing elongation off the end of the template (Fig 1C). In contrast to RNApII, TFIID remains associated under both conditions, showing its independence from the EC.

As discussed above, footprinting assays show TAFs interact with the TSS and regions downstream. A series of shorter immobilized templates showed 90 nucleotides downstream of the TSS supported post-transcription TAF binding, while 16 was insufficient (Fig 1D). Altogether, our biochemical data suggest a postinitiation conformation of TFIID bound to downstream DNA that does not require a TATA box for maintenance.

### In vivo evidence for downstream TAF binding

To determine the physiological relevance of downstream TAF binding, we looked for correlation between TAF occupancy and TAF dependence. TAF occupancy was defined as the number of Taf1 ChIP-exo reads (Rhee and Pugh, 2012) from -110 to +290 relative to the TSS. TAF dependence was defined as the difference in expression in two different *taf1* temperature sensitive alleles (**Fig S2A**) versus *TAF1* (Huisinga and Pugh, 2004). The data showed ribosomal protein genes (RPGs) were well separated from the other mRNA genes (**Fig 2A**). While RPGs have very high TAF occupancy and strong TAF dependence, there was surprisingly little correlation for the other mRNA genes. Therefore, we instead used standard deviation from the mean to put genes into four categories by their reliance on TAFs: RPGs, high (top 825 genes), medium (middle 4327), or low (bottom 729). Note that these categories are not meant to imply clearly discrete groups, but only to represent extremes along a spectrum.

In vivo evidence for differential downstream promoter interactions of TAFs came from re-examining high resolution ChIP-exo data from Rhee and Pugh (2012). They sorted genes by Taf1 occupancy and observed that low occupancy genes had Taf1 crosslinking patterns similar to the narrow peak seen for TBP and TFIIB upstream of the TSS. In contrast, high occupancy genes showed additional crosslinking overlapping the +1 nucleosome. We repeated this analysis using the four categories defined above. In plots of averaged values, Taf1 crosslinks appear to have three peaks: one overlapping TBP and TFIIB, one near the TSS, and one overlapping the +1 nucleosome (Fig 2B, C). Downstream peaks are also seen for Taf2 and Bdf1, but not several other TAFs (**Fig S2B**). Notably, Drosophila Taf2 also crosslinks predominantly downstream of the TSS (Shao and Zeitlinger, 2017). These results are consistent with a recent structural model of TFIID showing a downstream lobe consisting of Taf1, Taf2, and Taf7 (**Fig S2B**; Louder et al., 2016).

**Figure 2.**
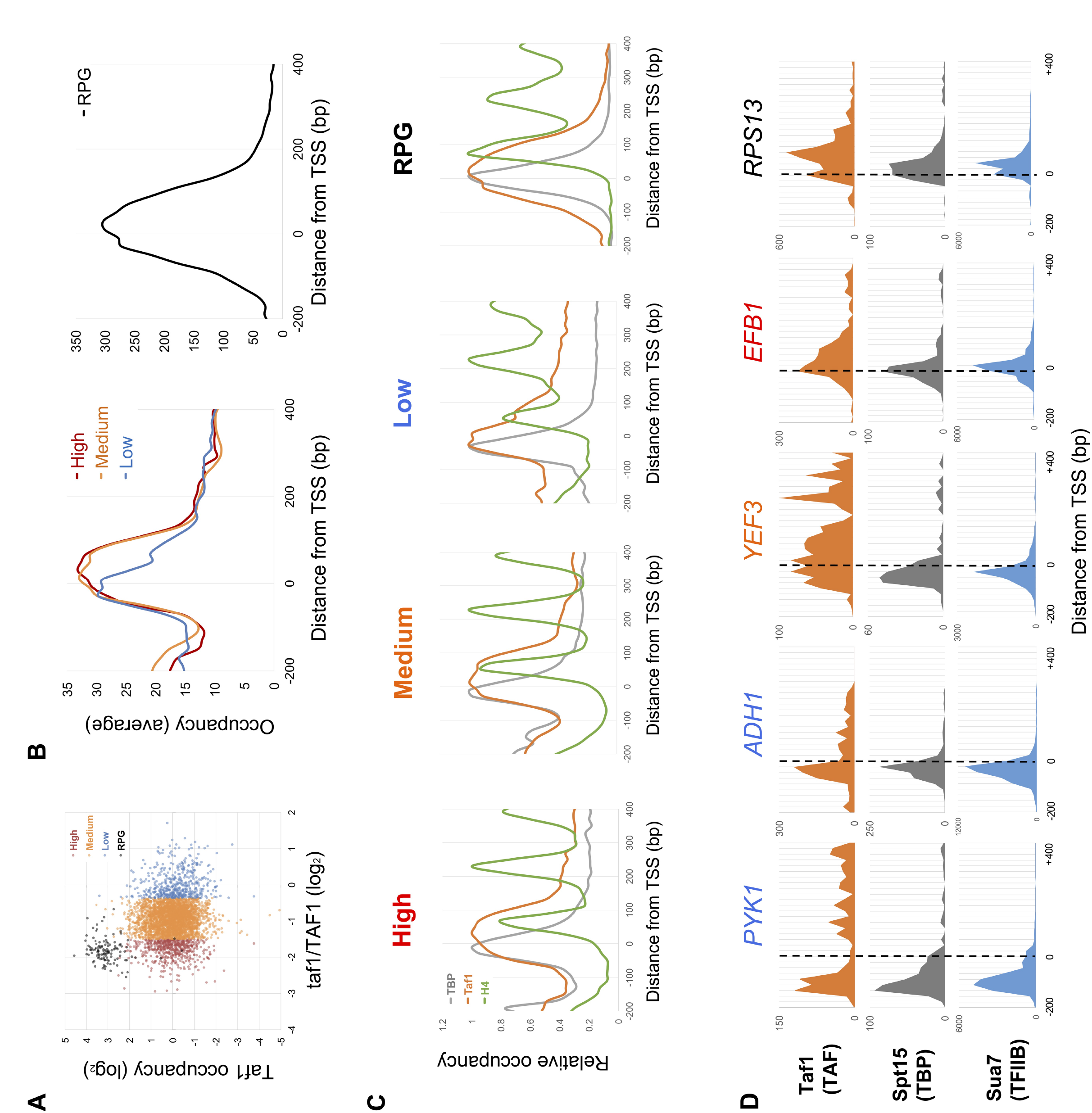
Taf1 downstream binding correlates with its transcription function. A. Scatter plot of Taf1 transcription dependence versus total occupancy. For all 4348 mRNA genes, the *taf1*/*TAF1* mRNA expression level ratios from (Huisinga and Pugh, 2004) were plotted in log2 scale on the x-axis. For each promoter, total Taf1 occupancy was plotted by summing ChIP-exo counts from -100 bp to +300 bp relative to the TSS (Rhee and Pugh, 2012), dividing by the average for all mRNA gene promoters, and plotting in log2 scale on the y-axis. Excluding the 137 RPGs, high TAF dependence genes (825) were defined as those with *taf1*/*TAF1* expression ratios below -1 standard deviation (SD = 0.57) from the mean of all gene ratios (0.95). Medium dependence genes (4327) had ratios between -1 and +1 SD, and low dependence genes (729) genes more than +1 SD from the mean of ratios. Average ratios in High, Medium, Low, and RPG groups are -1.73, -0.92, -0.08, and -1.82, respectively. See also **Fig S2A**. B. Averaged Taf1 occupancy for the four groups was plotted relative to the TSS (High: red, Medium: orange, Low: blue, and RPG: black) using ChIP-exo data from (Rhee and Pugh, 2012). See also **Fig S2B, C**. C. Occupancies for TBP, Taf1, and nucleosomes were plotted for the four groups separately to compare peak positions: TBP: gray, Taf1: orange, and histone H4: green. For each curve, data was normalized by setting maximum value to 1.0. D. Individual gene examples of Taf1 (TAF), Spt15 (TBP), and Sua7 (TFIIB) profiles at *PYK1* (Low *TAF1*-dependence: 0.02), *ADH1* (Low TAF1-dependence: 0.35), *YEF3* (Medium *TAF1*-dependence: -1.21), *EFB1* (High *TAF1*-dependence: -2.02), and *RPS13* (RPG, *TAF1*-dependence: -1.00).

Average values of Taf1 overlapping the TBP peak are roughly equal in high, medium, or low TAF-dependence classes (Fig 2B). However, promoters with medium or high TAF-dependence show highest levels of Taf1 crosslinking near and downstream of the TSS in averaged plots (Fig 2C), as well as when ratios of downstream to upstream crosslinking are calculated for individual genes (**Fig S2C)**. Although RPG promoters have roughly ten-fold higher overall Taf1 crosslinking, their normalized profile is most similar to the high TAF-dependence promoters. These trends can also be seen in individual gene traces (Fig 2D). Therefore, the response of promoters to *TAF1* mutations is not correlated with absolute level of Taf1 binding, but instead with downstream promoter interactions of TFIID.

### Downstream TFIID binding correlates with Bdf1 and acetylated histone H4

We previously isolated Bdf1 as a Taf7-interacting protein corresponding to the BET module of mammalian Taf1 (Matangkasombut et al., 2000). Bdf1 was also identified as a subunit of the SwrC complex, which replaces H2A with H2A.Z/Htz1 (Kobor et al., 2004; Krogan et al., 2003; Mizuguchi et al., 2004). The Bdf1 bromodomains interact with acetylated H4 tails (Ladurner et al., 2003; Matangkasombut and Buratowski, 2003). As histone H4 acetylation (H4ac) is highest at +1 nucleosomes of active promoters, Bdf1 and other BET proteins are thought to recruit their associated complexes to TSS regions.

Since Bdf1 behaved like TAFs, but not SwrC, in our immobilized template system, we asked which complex its behavior most closely resembled in vivo. ChIP-seq for TBP, Taf1, histone H3, and H4ac was performed. Consistent with their high transcription rate, RPG promoters have lowest H3 occupancy but highest H4ac (**Fig S3A**). Average H3 levels were similar in other gene classes, but low TAF dependence genes had notably less H4ac than high or low dependence genes. These results were aligned (Fig 3A) with previously published datasets for Bdf1, Swr1, and Htz1 (Gu et al., 2015; Rhee and Pugh, 2012; Yen et al., 2013). Genes were sorted into the four classes of TAF-dependence and then further within each category by the Taf1 downstream crosslinking signal from +30 to +170 relative to the TSS. Bdf1 correlated with Taf1 and TBP better than Swr1 and Htz1 (**Fig S3B**). Taf1 and Bdf1 were particularly high at RPGs, while Swr1 and Htz1 were largely absent (Fig 3A, **S3B**). H4ac correlated best with Bdf1, with both Swr1 and Taf1 also showing some correlation. Importantly, TBP correlated well with Taf1 and Bdf1 on TAF-dependent genes, while genes with low TAF dependence had high TBP levels even when Taf1 levels were low (Fig 3A, **S3B**).

**Figure 3.**
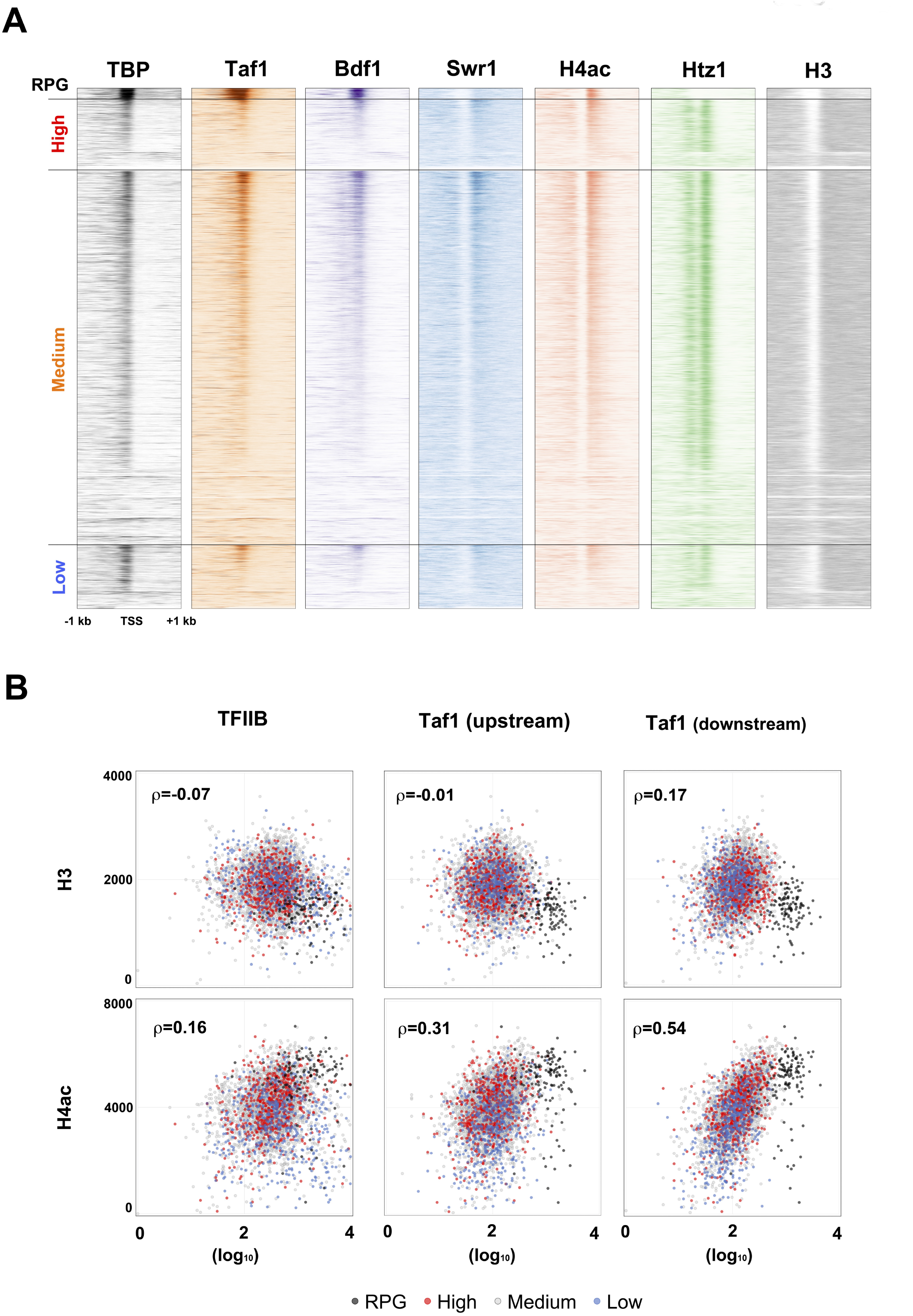
Downstream Taf1 binding correlates with +1 nucleosome H4 acetylation. A. ChIP patterns of TBP, Taf1, Bdf1, Swr1, H4ac, Htz1, and H3 at individual genes were plotted as horizontal lines to generate heatmaps, with darker color signifying higher levels. Genes were sorted into the four *TAF1*-dependence groups, and then within each group by Taf1 downstream occupancy (ChIP-exo reads from +20 bp to +160 bp relative to TSS). Together with our newly generated data, raw data for Bdf1 (Rhee and Pugh, 2012), Swr1 (Yen et al., 2013), and Htz1 (Gu et al., 2015) were reanalyzed as in Materials and Methods. See also **Fig S3A**. B. H3 and H4ac levels were plotted versus TFIIB and Taf1 binding. Occupancies of H3 and H4ac were determined as the sum of ChIPseq reads from +30 bp to +170 bp relative to TSS, encompassing the +1 nucleosome. Occupancies of Taf1 and TFIIB were determined as total ChIPexo reads (Rhee and Pugh, 2012) from -140 bp to -1 bp for the upstream core promoter and +20 bp to +160 bp for the downstream promoter. Each dot represents one mRNA gene (total 4720). For each graph, Spearman’s rank correlation coefficient (ρ) is shown. See also **Fig S3B**.

Taf7 binds to both downstream Taf1/Taf2 (Louder et al., 2016) and Bdf1 (Matangkasombut et al., 2000), presumably positioning the Bdf1 bromodomains similarly to those found in metazoan Taf1. If so, bromodomain interactions with H4ac could promote the downstream DNA binding conformation of TFIID. In agreement with this prediction, H4ac correlates better with Taf1 crosslinking to downstream promoter regions (+30 to +170) than to upstream Taf1 (-150 to -10) or the PIC as assayed by TFIIB signal (Fig 3B, **S3B**). These results are consistent with a model where Bdf1 (or the metazoan Taf1 bromodomains) bind H4ac to help tether TFIID to the +1 nucleosome and downstream promoter region.

### TAFs and transcription promote activator-independent transcription

One likely function for stable post-transcription interaction of TAFs with downstream promoter DNA is to promote reinitiation. To test this idea, nuclear extracts were prepared from isogenic wild-type (*TAF1*) or temperature sensitive *taf1* strains (*taf145 ts-1* strain in Walker et al., 1996). Single round transcription in the *taf1* mutant extract was similar to or better than wild type, and both extracts showed a 2- to 3-fold increase in the presence of the transcription activator Gal4-vp16 (Fig 4A, **S4**). Increased transcript levels were similarly produced by multiround conditions in the presence of activator. In contrast, the *taf1* extract only produced about half as much transcription in the multi-round reaction in the absence of activator (Fig 4B, **C, S4**).

**Figure 4.**
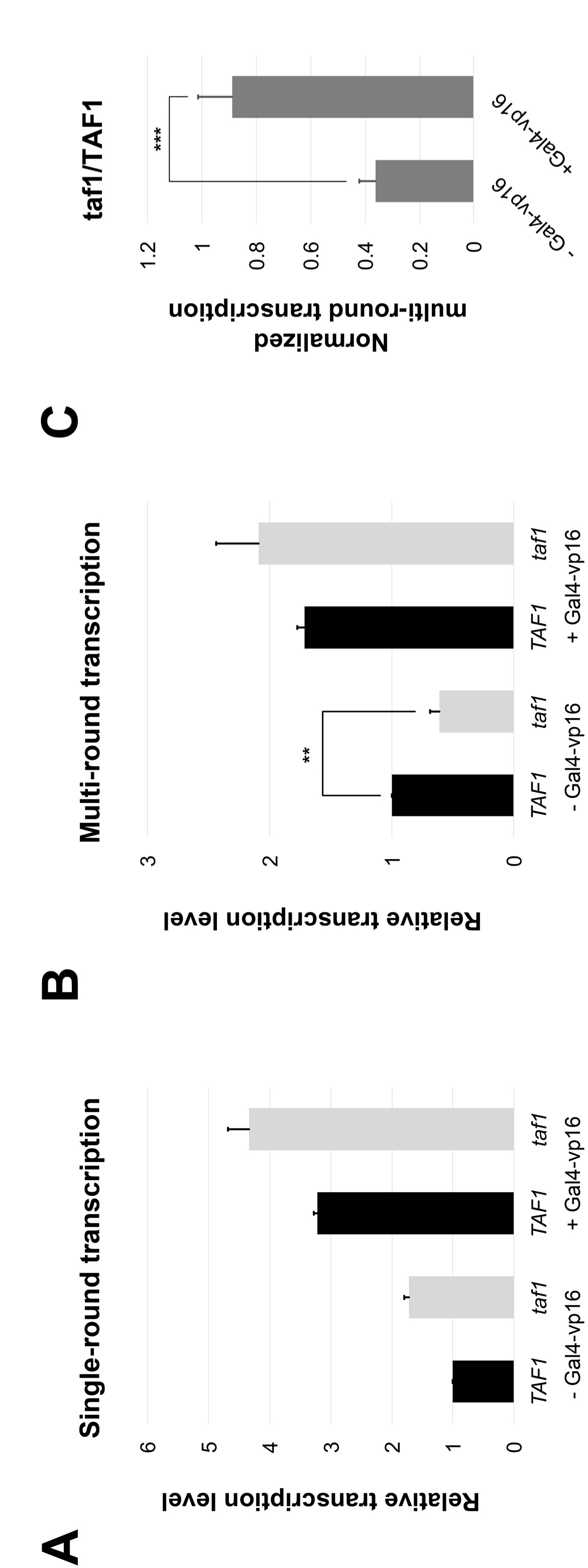
Taf1 promotes activator-independent multi-round transcription. *In vitro* transcription was performed with *TAF1* (Black, YF157/YSW87) or *taf1* (Gray, YF158/YSW90) yeast nuclear extracts. NTPs were added for (A) 3 minutes insingle-round reactions, or (B) 45 minutes for multi-round transcription, in the presence or absence of transcription activator Gal4-vp16 as indicated. Transcripts were quantified, normalized, and plotted as the average value from three independent reactions. Error bars show standard deviation (**: p = 0.014, ***: p=0.006; unmarked pairs p > 0.01). See also **Fig S4**.

To test if TAF-dependent stimulation of multi-round transcription is connected to the transcription-dependent TAF deposition described above, a two-step reaction scheme was used (Fig 5A). In the first step, nuclear extract is mixed with Template 1 immobilized on magnetic beads, exactly as in the proteomics experiments in Fig 1. Pre-incubation with unlabeled NTPs allows transcription, or instead transcription can be blocked by omitting NTPs or by adding α-amanitin with the NTPs. The complexes associated with Template 1 are then washed extensively and added to a second reaction containing nuclear extract, a second Template 2 as an internal control for de novo initiation, and radioactive NTPs for quantitation of transcription from both templates. Stimulation of reinitiation manifests as faster accumulation of transcripts from Template 1 relative to Template 2. As re-initiations also occur on Template 2 after the initial round, the increase will not be cumulative over time and will have a maximum value of 2-fold (i.e. if every Template 1 reinitiates once before any Template 2 initiation in the second reaction).

**Figure 5.**
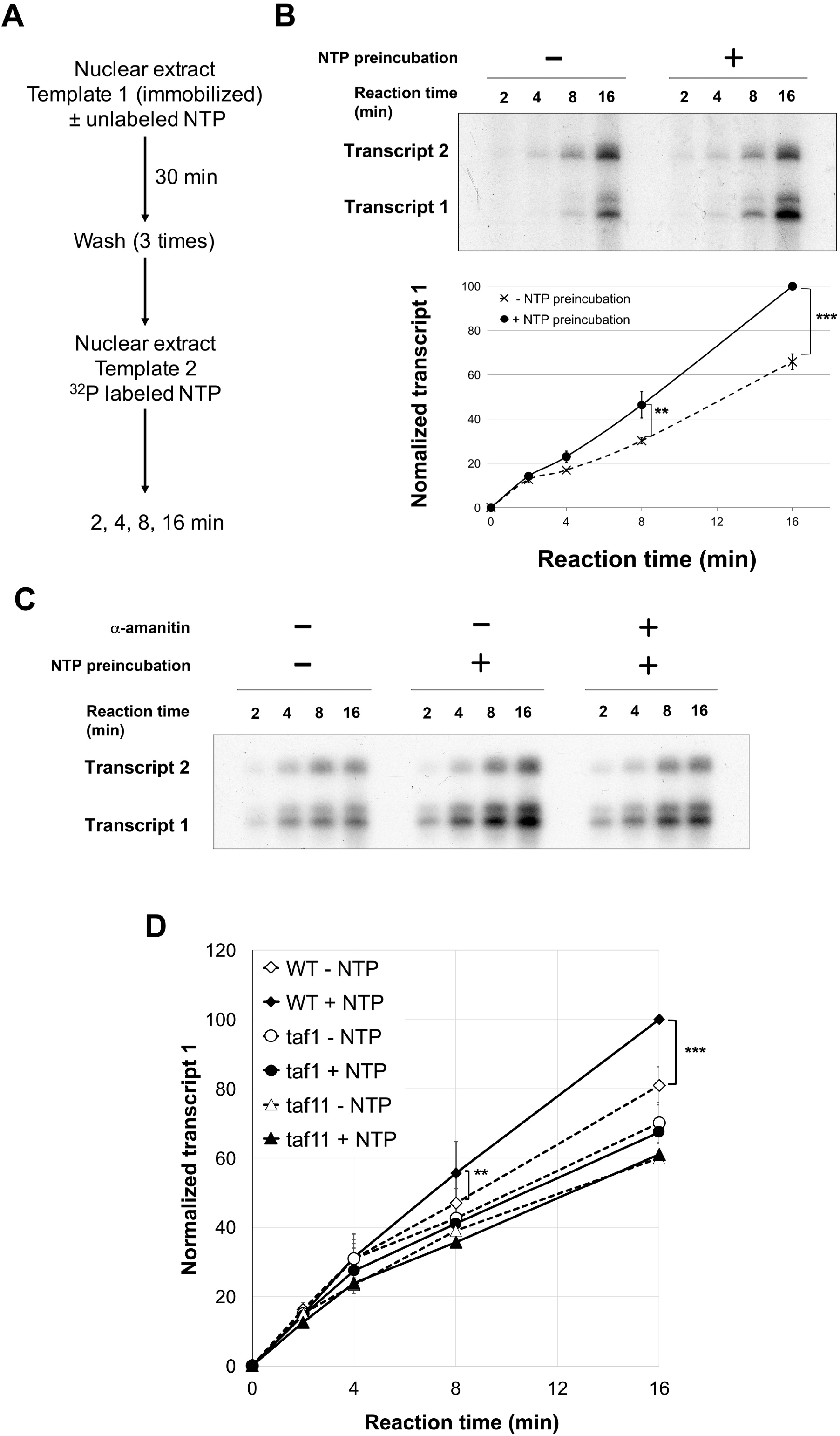
Taf1 promotes transcription reinitiation. A. Schematic of two-step sequential reactions used to analyze transcription reinitiation in vitro. Note that no transcription activator is used in these experiments. B. Yeast nuclear extract was pre-incubated with immobilized Template 1 (as in Fig 1A) in the presence or absence of NTPs (400 μM each). After three washes, nuclear extract, ^32^P-labeled NTPs, and the longer G-less cassette Template 2 (pG5CG-D2/F916) were added. After 2, 4, 8, and 16 min, labeled transcripts were isolated and analyzed by gel electrophoresis and autoradiography (upper panel). Quantitation from three independent reactions (bottom panel) plots Transcript 1 levels over time, normalized by setting the 16 minute maximum to 100 and using Transcript 2 to correct for variations in recovery or reaction conditions. Error bars indicate standard deviation. See also **Fig S5A**. P value: ** = 0.01, *** = 7.5 x 10^−5^. C. The sequential transcription assay was performed with NTPs and α-amanitin (10 μg/ml) in the pre-incubation as indicated. D. Sequential transcription assay comparing wild-type (YSW87) to mutant *taf1* (YSW90) or *taf11* (YSB1732) nuclear extract in the first pre-incubation reaction. After washes, the second transcription reaction was performed with wild type nuclear extract prepared from strain BJ2168. Quantitation from three (*taf11*) or five (*taf1*) independent replicates is shown. P value: ** = 0.08, *** = 5.2 x 10^−5^. All other differences were not statistically significant. See also **Fig S5B**.

When wild type extract was used for both steps, a marked increase in Template 1 transcription was seen when NTPs were included in the first reaction (Fig 5B). Stimulation was strongly decreased by inclusion of α-amanitin in the first reaction (Fig 5C), arguing that transcription per se, and not some other NTP function, mediates the effect. Interestingly, the reinitiation effect was not seen when Gal4-vp16 was included in the reaction (**Fig S5A**), possibly because a high level of activator-induced de novo initiation swamps the reinitiation signal. Finally, when *taf1* or *taf11* mutant extract is used for the first reaction, no stimulation of reinitiation was observed, even though wild-type extract was used for the second reaction (Fig 5D, **Fig S5B**).

Together, our in vitro results reveal the existence of a reinitiation intermediate that is transcription-dependent, TAF-dependent, and activator-independent. While superficially similar to the “Scaffold” complex defined by Yudkovsky et al., (2000), multiple lines of evidence make the differences clear (see Discussion). These reinitiation intermediates are not mutually exclusive, and the residual stimulation we observe with α-amanitin in the first reaction (Fig 5C) could represent Scaffold-stimulated reinitiation.

### In vivo evidence for two TFIID states

Based on the observations above, we developed the following model (also see Discussion below). First, a transcription activator promotes a pioneer round of transcription by recruiting coactivators such as SAGA and Swi/Snf to move promoter-occluding nucleosomes. A PIC then assembles, stimulated by activator-Mediator interactions. In this initial round of transcription, TAFs may primarily be tethered upstream to TBP or even be absent from the initial PIC. After initiation and promoter clearance by RNApII, the TAFs are can then bind downstream, where they could promote one or more cycles of reinitiation. This model predicts at least two states of TFIID, distinguished by whether the Taf1/Taf2/Taf7/Bdf1 module is bound to the downstream promoter or not.

This idea was tested using Anchor Away depletion (Haruki et al., 2008) of Rpb1 or Bdf1 from nuclei. ChIP-seq of Taf1 and TBP was performed before and after a rapid 60 minute depletion of the indicated factor. The relative change was plotted by heat map of individual promoters (Fig 6A), or by gene averaged plots (**Fig S6**). Depletion of Rpb1 led to a widespread drop in TBP crosslinking, presumably because lack of RNApII limits PIC formation. Taf1 also dropped throughout the entire length of RPG and low TAF-dependence promoters, but behaved quite differently on high and medium promoters (Fig 6B). Within this group, despite the drop in TBP, Taf1 crosslinking decreased upstream while at the same time increasing downstream. This shift is consistent with a TAF complex binding downstream of the TSS after departure of RNApII.

**Figure 6.**
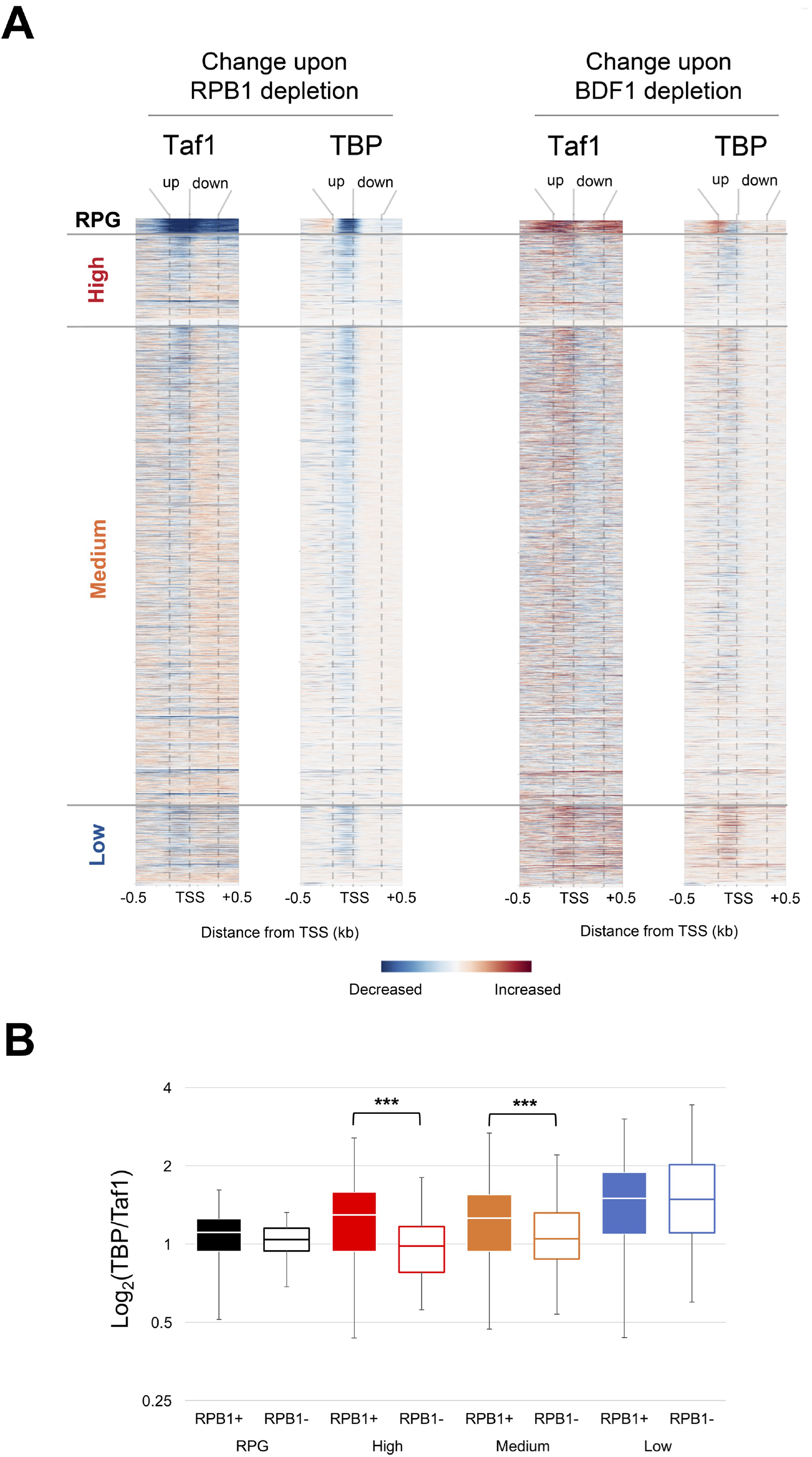
Depletion of Rpb1 or Bdf1 alters Taf1 binding in vivo. A. ChIPseq for Taf1 or TBP Rpb1 was performed from cells before or after a 60 minute depletion of Rpb1 or Bdf1 by Anchor Away. Normalized reads (reads per million) were mapped and differences plotted as heat maps, showing individual genes sorted by the classes defined in Fig 2. See also **Fig S6A, B**. B. The ratio of total TBP and Taf1 reads for each promoter were calculated and plotted as quartile box and whisker plots.

At promoters with high or medium TAF dependence, Bdf1 depletion had the opposite effect as Rpb1 depletion on Taf1 binding. Supporting a role for Bdf1 in anchoring TFIID downstream, Taf1 crosslinking decreased downstream while remaining the same or increasing upstream. TBP was unaffected or slightly decreased. In contrast, TBP and Taf1 in both regions increased on low TAF-dependence promoters, perhaps due to redistribution of TFIID from the other promoters. Taf1 also increased on RPG promoters, although less so in the region just downstream of the TSS. While unexpected, it should be noted that cells lacking Bdf1 grow very slowly yet are viable due to the presence of its homolog Bdf2. The Bdf2 bromodomains do not show preference for acetylated H4 (Matangkasombut and Buratowski, 2003), so depletion of Bdf1 may result in re-targeting of TFIID by Bdf2.

In summary, the differential response of Taf1 crosslinking to regions upstream and downstream of the TSS is consistent with independent binding of TFIID modules. Bdf1 promotes downstream interactions while RNApII antagonizes them.

## DISCUSSION

In the process of isolating and characterizing RNApII elongation complexes, we unexpectedly discovered that transcription initiation results in stable association of TAFs with downstream promoter regions, independent of TBP bound to the TATA box. Transcription-dependent deposition of a TAF complex is reminiscent of the Exon Junction Complex, which is deposited just upstream of splice junctions by the spliceosome as a marker of recent intron removal (Moore and Proudfoot, 2009). The downstream TAF complex similarly creates a "memory" of recent transcription, promoting activator-independent re-initiation. Incorporating our data with earlier observations, we propose the following model (Fig 7A).

**Figure 7.**
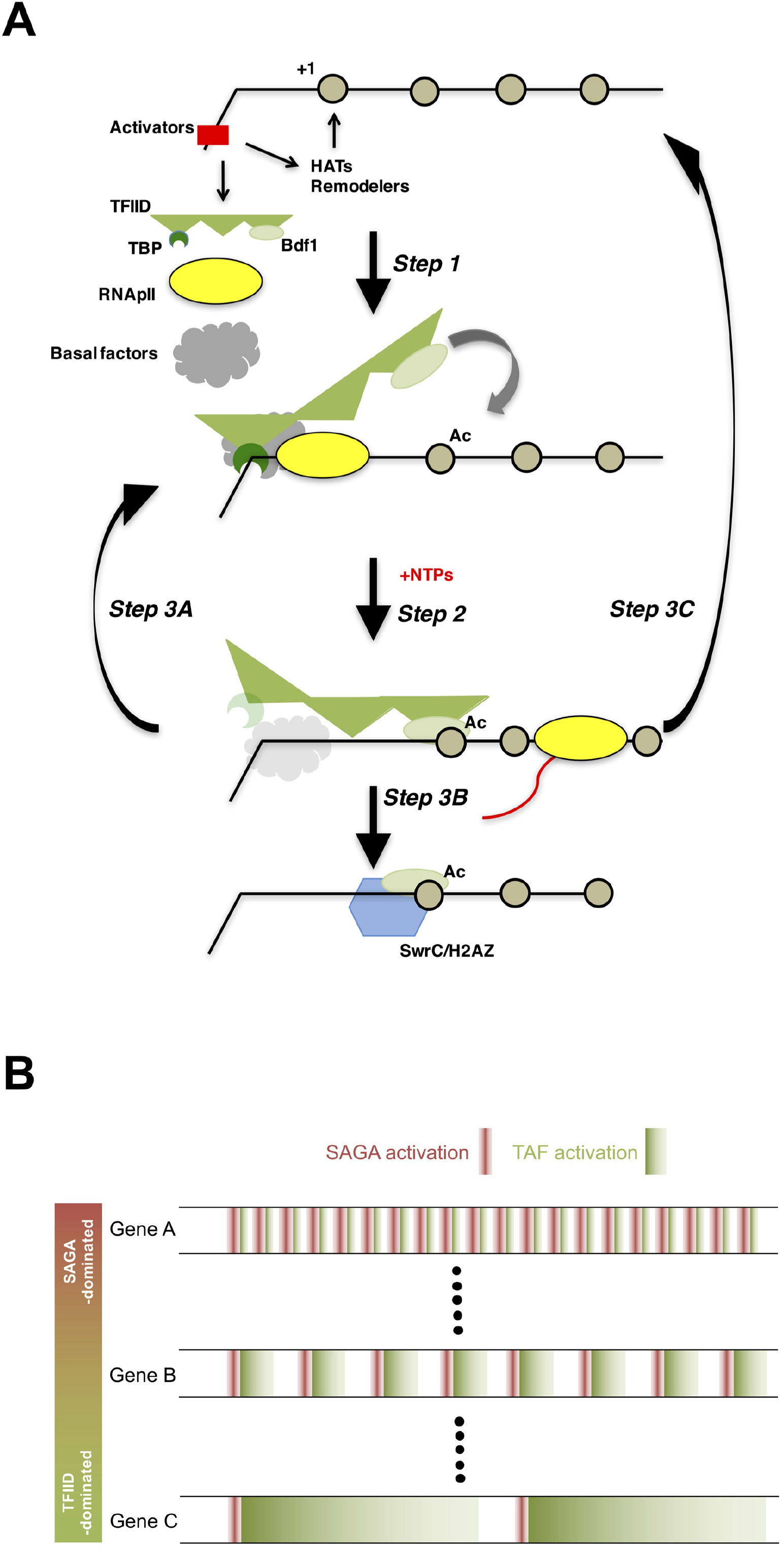
Mechanism of TAF-mediated RNApII transcription. A. Model for how TAFs may promote reinitiation. See Discussion for details. TFIID is shown as TBP (dark green), TAFs (three flexibly linked triangles in medium green), and Bdf1 (light green oval). DNA is represented as a black line, nucleosomes as beige circles, transcription activators as a red rectangle, basal factors as a gray cloud, RNApII as a yellow oval, SwrC as a blue hexagon, and the RNA transcript as a red line. The lighter versions of TBP and basal factors after Step 2 is to signify that these factors may be largely or completely dissociated after initiation. B. A reinitiation model for how two promoters of similar strength can exhibit differential responses to TAF or activator/SAGA mutants (Huisinga and Pugh, 2004; de Jonge et al, 2016). Each horizontal trace represents behavior over time. An initial activator- and SAGA-stimulated round of transcription is represented in red, while subsequent TAF-stimulated reinitiations are represented in green.

### Step 1: Pioneer round of PIC assembly

Regulatory transcription factors recruit HATs (SAGA and NuA4 in yeast) and ATP-dependent chromatin remodelers to move occluding nucleosomes from the core promoter (Cairns, 2009; Weake and Workman, 2010). Several mechanisms for TBP delivery have been described, and which is used may depend on which activators are present at the promoter. It has been proposed that TBP can arrive at the promoter via its association with SAGA (Eisenmann et al., 1992; Mohibullah and Hahn, 2008), as free TBP (Buratowski et al., 1988), or as part of TFIID. If the latter, TFIID may be recruited by specific promoter sequences (Kadonaga, 2012) or activators (Albright and Tjian, 2000; Chen et al., 2013; Mencía et al., 2002; Papai et al., 2010). Once TBP in place, the other basal factors, including Mediator-RNApII, assemble into the PIC.

### Step 2: Initiation and establishment of downstream TAF interactions

NTPs trigger promoter melting and synthesis of RNA. Upon promoter escape by RNApII, some factors may remain associated to facilitate reinitiation. An early in vitro study with purified mammalian basal factors found only TFIID remaining on immobilized templates after transcription (Zawel et al., 1995). Yudkovsky et al. (2000) described a yeast "Scaffold" complex containing TFIID, TFIIA, Mediator, TFIIH, and possibly TFIIE persisting at the promoter. Recent ChIP-seq and ChEC-seq experiments argue that Mediator primarily resides at the activator binding site and only transiently at the basal promoter (Grünberg et al., 2012; Jeronimo and Robert, 2014; Wong et al., 2014), so Scaffold may actually be two complexes: an activator-Mediator complex at the UAS and a subset of basal factors at the core promoter.

Our data suggest an alternative reinitiation intermediate in which TAFs interact with downstream promoter DNA and the acetylated +1 nucleosome. This complex is clearly distinct from Scaffold. First, Scaffolds were defined as what remains after extensively washed PICs were treated with NTPs or ATP only. In contrast, our initial round of transcription occurs in extract, allowing post-initiation recruitment, as seen with TAFs (**Fig S1D**). Whereas Scaffold is dependent on activator, our reinitiation effect does not require activator (Figs 4, **S4A**). Finally, while Scaffold can be generated with ATP only, our reinitiation intermediate requires actual transcription (Figs 1, 5C).

In vivo evidence for downstream interactions in our reinitiation intermediate comes from patterns of Taf1 crosslinking (Figs 2, 3). At TFIID-dominant promoters, downstream crosslinking increases after rapid RNApII depletion, even as upstream crosslinking drops (Fig 6A). Interestingly, a recent ChEC-Seq experiment from Grünberg et al (2016) showed that depletion of Mediator subunit Med14 caused increased Taf1-mediated cleavage downstream of the TSS. Other groups showed that TBP inactivation unexpectedly increased TAF crosslinking at some promoters (Mencía et al., 2002; Shen et al., 2003), Originally interpreted as TBP-independent binding of TAFs, our model suggests these TAFs instead remain from previous rounds of transcription.

Transcription-dependent TAF deposition was seen in vitro on both naked and chromatinized templates (Fig 1, **S1E**). However, interactions between the Taf1/Bdf1 bromodomains and the acetylated +1 nucleosome are also likely important for TFIID function in vivo. A recent TFIID structure lacks the bromodomains, but it shows a Taf1/Taf7 module binding the promoter 50-60 base pairs downstream of the TATA box (Louder et al., 2016). Bdf1 specifically interacts with Taf7 (Matangkasombut et al., 2000), consistent with Taf1/Bdf1 bromodomains mediating communication between the +1 nucleosome and TFIID. Supporting this idea, acetylated H4 correlates better with downstream than upstream Taf1 crosslinking (Fig 3B, **S3B**). Even more compellingly, depletion of Bdf1 results in a reduction of Taf1 crosslinking specifically downstream of the TSS (Fig 6B).

### Step 3: A TAF-mediated branchpoint

Depending on the particular promoter sequences and chromatin modifications, three outcomes can be envisioned: A) assembly of a new PIC for reinitiation, B) dissociation of TAFs but not Bdf1, leading to SwrC recruitment and H2A.Z incorporation at the +1 nucleosome, or C) TFIID dissociation and reversion to the nucleosome-occluded state.

### Option 3A: TAF-dependent reinitiation

We propose the TAF complex stimulates reinitiation through two mechanisms. First, it can sterically block ingression of the +1 nucleosome to an occluding position, bypassing the need for activator-dependent recruitment of HATs and remodelers as in Step 1. Second, the TAF complex can stabilize TBP or perhaps recruit a new TBP molecule. This might be particularly significant at promoters with weak TATA boxes.

At SAGA-dominant but not TFIID-dominant promoters, the +1 nucleosome overlaps the PIC (Rhee and Pugh, 2012). Two recent papers show that the +1 nucleosome markedly shifts downstream when promoters with low TAF occupancy become active (Nocetti and Whitehouse, 2016, Zhou et al, 2016). In contrast, promoters that are highly expressed or with high TAF occupancy have a more static +1 nucleosome at the downstream-shifted position. Finally, Reja et al. (2015) showed that transcription inhibition causes the +1 nucleosome to shift upstream over TFIID-dominant RPG promoters. Together, these observations support a model where TAFs help maintain the +1 nucleosome at a downstream position.

As TAF-dominant promoters tend to have weaker, non-consensus TATA boxes, downstream TAF interactions likely also help retain or rebind TBP at the TATA box. Supporting a retention model, an in vivo epitope-switching experiment (van Werven et al., 2009) found highest TBP turnover at SAGA-dominant promoters, and lowest TBP turnover at promoters with high Taf1 levels.

Importantly, TAF mutant extracts support single-round transcription as efficiently as wild-type extract, but have much less activity in multi-round reactions (Fig 4, 5). Similarly, TAF depletion from mammalian extracts did not affect activator-induced transcription in single-round transcription reactions, but strongly reduced transcription in multi-round reactions (Oelgeschläger et al., 1998). Whether the post-initiation TFIID complex assembles a new PIC by reusing basal factors from the Scaffold or recruiting them de novo, this new cycle of transcription bypasses the need for the activator-dependent Step 1.

### Option 3B: Recruitment of SwrC and H2A.Z

If TAFs dissociate from downstream DNA before a new PIC can assemble, it is interesting to speculate that, in yeast, the acetylated +1 nucleosome might retain Bdf1. This could create a window for Bdf1 to recruit SwrC complex for incorporation of H2A.Z/Htz1 at the +1 nucleosome. Yeast Htz1 levels are highest at inactive promoters with acetylated +1 nucleosomes (Li et al., 2005; Zhang et al., 2005) (Altaf et al., 2010). It was proposed that Htz1 binding provides a "memory" of recent transcription to allow rapid gene reactivation. By our model, Htz1 incorporation would occur most frequently at genes with relatively high levels of both de novo initiation (Step 1) and TAF-dependent reinitiation (Step 3A). In agreement, genome-wide analysis shows genes with highest Swr1 and Htz1 levels are most likely to have medium TAF dependence (Fig 3A). It will be informative to monitor the dynamics of histones, TAFs, and SwrC at these promoters.

### Option 3C: Return to the inactive state

TFIID dissociation could allow the promoter to revert to the repressed state at the beginning of Step 1. The lifetime of post-initiation TFIID at promoters may vary depending on particular DNA sequence elements, TAF-recruiting activators, and +1 nucleosome acetylation.

The relative probabilities of TAF-dependent reinitiation versus returning to the repressed state provides a simple explanation for the spectrum of responses to SAGA and TAF mutations, without having to postulate completely distinct initiation mechanisms (Fig 7B). Promoters with weak TAF retention will rarely reinitiate, and the need to continuously acetylate and remodel the +1 nucleosome will create strong SAGA-dependence. In contrast, those with strongest TAF retention will have more reinitation relative to the initial round of transcription and be most TFIID-dominant. It is likely these behaviors also contribute to different "transcription burst" properties of promoters. Transcription bursting is most common at SAGA-dominant promoters with strong TATA boxes, while TAF-dominant promoters generally have lower levels of noise (Blake et al., 2006; Raser and O’Shea, 2004).

One prediction of our model is that SAGA-dominant promoters will have a stronger requirement for the continued presence of activator. In striking agreement, de Jonge et al. (2016) recently showed that nuclear depletion of the Hsf1 transcription activator causes rapid loss of transcription at its SAGA-dominant, but not its TFIID-dominant, target promoters. There is a growing consensus that the SAGA and TFIID pathways are not mutually exclusive, but instead work redundantly or cooperatively at most promoters (Bonnet et al., 2014; Grünberg et al., 2016)(Huisinga and Pugh, 2004). Our model accommodates this fact while explaining the differential responses of promoters to SAGA and TFIID mutations.

### TFIID conformations and transcription-dependent TAF interactions

An important future goal will be to determine the physical basis for transcription-dependent TAF binding. A recent partial model for TFIID bound to DNA (Louder et al., 2016) reveals some of the downstream promoter interactions likely to be relevant. This structure can be docked into a separate model of the PIC built on TBP, so it may be feasible to accommodate all these proteins simultaneously. However, careful kinetic experiments by Yakovchuk et al. (2010) indicate that TAFs actually retard PIC assembly, and that TFIID downstream promoter contacts rearrange or release as RNApII forms a closed complex. TAF downstream DNA contacts are almost certainly released at some point in the transcription cycle, at the very least during early transcription when downstream DNA is drawn into the RNApII active site by TFIIH (Hahn and Buratowski, 2016).

At this point, TAFs may dissociate, only to be recruited again at a later step during promoter clearance or early elongation. In a more likely model, a conformation change may move TAFs on and off the DNA while retaining contact with TBP or other factors. This may simply reflect competition between TAFs and RNApII for downstream regions, but it also possible that some event or modification during promoter clearance actively places TAFs downstream. The different responses of upstream and downstream Taf1 crosslinking upon Rpb1 or Bdf1 depletion support the concept of at least two TFIID conformations (Fig 6). Flexibility of TFIID is clear from early footprinting experiments (Chi and Carey, 1996; Horikoshi et al., 1988a; 1988b) as well as recent cryo-EM studies (Cianfrocco et al., 2013; Elmlund et al., 2009; Papai et al., 2010). Notably, glutaraldehyde fixing was needed to stabilize the Louder et al. (2016) TFIID structure with downstream binding.

Previous models presumed that TBP and TAFs act in concert as a single promoter recognition factor. Our model suggests TBP and TAF binding to DNA can occur more independently at different stages of the transcription cycle. Interestingly, based on comparisons of free and DNA-bound TFIID structures, Nogales and colleagues speculated there could be a TFIID conformation having downstream TAF-DNA contacts while TBP and upstream TAFs release from the TATA-box (Cianfrocco et al., 2013). Such a complex is consistent with the reinitiation intermediate suggested by our results. The ability of a TBP-TAF complex to independently release upstream or downstream contacts may also resemble events during RNA polymerase III transcription, where TFIIIB and TFIIIC subunits interact with promoter elements downstream of the transcription start site to assemble the PIC, yet somehow do not interfere with elongation. The TAF reinitiation function described here helps explain some previously confusing TFIID results and provides a new model for thinking about TFIID functions.

## METHODS

### Strains, plasmids and oligonucleotides

*S. cerevisiae* strains, oligonucleotides and plasmids used in this study are listed in **Table S2, S3, S4**.

### Yeast nuclear extract preparation

Nuclear extracts were prepared as described previously (Sikorski et al., 2012). Wild type extracts for mass spectrometry analysis were prepared from strains BY4741 or BJ2168/YF4. For in vitro transcription assays, nuclear extracts were prepared from wild-type (YSW87/YF157), *taf1 ts-1* (YSW93/YF158), or *taf11-3000* (YSB1732) strains (Walker et al., 1996; Komarnitsky, Michel, and Buratowski, 1999).

### Immobilized template binding and quantitative mass spectrometry

Immobilized template binding assay was performed as described (Sikorski et al., 2012) with some modifications. Biotinylated linear templates were amplified by PCR from pUC18-G5CYC1 G-(SB649) with primers listed in **Table S3**. Where noted, templates were chromatinized with yeast histone octamers as described (Sikorski et al., 2012). Chromatinized or naked templates (880 ng DNA) were immobilized by binding to 200 μg streptavidin beads (Dynabeads Streptavidin T1, Invitrogen) for one hour at room temperature (RT). Beads were isolated by magnetic concentration and then blocked in 600 μl blocking solution [1X transcription buffer (100 mM K-acetate, 20 mM HEPES-KOH pH 7.6, 1 mM EDTA and 5 mM Mg-acetate) with 60 mg/ml casein, 5 mg/ml polyvinylpyrrolidone, 2.5 mM dithiothreitol (DTT)] for 25 min at RT. Transcription activator Gal4-vp16 (800 ng) was incubated with templates for 5 min at RT. Yeast nuclear extracts (~1.5 mg) were added in 240 μl of 1X transcription buffer complemented with 80 units RNAsin, 0.44 units creatine phosphokinase, 307.2 μg phosphocreatine, 0.03% NP-40, 2 μg tRNA, 160 μM S-adenosylmethionine, 20 μM acetyl-CoA, and 4 μg HaeIII-digested *Escherichia coli* DNA for 30 min at RT. Transcription was started by adding NTP mix (400 μM each ATP, CTP, UTP, and 3’-OmeGTP) to the mixture. Where indicated, α-amanitin was added at 10 μg/ml. The reaction was stopped after 15 min by washing twice in 200 μl Wash buffer (1 X transcription buffer complemented with 0.05% NP-40 and 2.5 mM DTT). Downstream DNA and bound proteins were released with SspI-HF or PstI-HF digestion (120 units, New England Biolabs) in 160 μl 1 X transcription buffer for 30 min at RT. Sample preparation for quantitative mass spectrometry and subsequent data analysis were performed as described (Sikorski et al., 2012). For relative quantitation, iTRAQ reporter signal intensity values of all unique peptide scans for a given protein were summed prior to calculation of ratios. All iTRAQ ratios are normalized to total reporter signal. Only proteins that were identified in multiple replicates were used. Statistical significance of enrichment was calculated by t-test (**Fig S1A**) and Mixed Model (**Fig S1B, C**) analyses.

### Chromatin immunoprecipitation and ChIP-seq analysis

For Fig 3, strain BY4741 was used. For nuclear depletion of Rpb1 (YSB3202) or Bdf1 (YSB3323) in Fig 6, the Anchor-Away system (Haruki et al., 2008) was used, with 1 μg/ml rapamycin added for one hour to cells containing the appropriate FRB fusion. Cells were grown to OD_600_ = 0.5 and then crosslinked with 1% formaldehyde for 20 min at RT. Crosslinking was stopped with 125 mM glycine for 5 min, and cells were collected and washed. Sheared chromatin (100 bp ~ 300 bp) was prepared using a Misonix sonicator 300 with cup horns.

Crosslinked chromatin was precipitated by standard methods with specific antibodies against Taf1 (provided by Joe Reese, Penn State Univ.), Spt15 (TBP), H3 (Abcam, ab1791), or H4ac (Millipore, 06-598). Libraries for multiplex sequencing were generated as described previously (Wong et al., 2013) and sequenced on an Illumina 2500 machine by the Harvard Bauer Center Genomics core facility. Sequenced reads were aligned to the yeast S288C reference genome (www.yeastgenome.org, with annotations provided by Burak Alver, HMS) using Bowtie 1 (Langmead et al., 2009). MACS (version 2) was used to visualize peaks from sequence read distribution (Feng et al., 2012). Custom Python scripts for analyzing and graphing data (L. Soares, manuscript in preparation) are available at https://github.com/Buratowski/NGS. Data is deposited in GEO (accession number GSE94851). In order to reanalyze genome wide occupancy, raw data for Taf1, Bdf1 (Rhee and Pugh, 2012), Htz1 (Gu et al., 2015), and Swr1 (Yen et al., 2013) were adapted and reanalyzed in parallel as above (GEO accession numbers: Taf1: SRR396796, Bdf1: SRR397550, Htz1: SRR1118262, Swr1: SRR948403).

### *In vitro* Transcription

*In vitro* transcription was performed as described (Sikorski et al., 2012). After pre-incubation of template and extract, NTPs were added for three minutes for single-round transcription and for 45 minutes for multi-round reactions. For the two-step transcription analysis in Fig 5, NTP pre-incubation was performed as in the Immobilized Template Binding Assay as describe above, except Gal4-vp16 was not added. Template 1, shown in Fig 1A, was incubated with wild-type, *taf1*, or *taf11* mutant extract and unlabeled NTPs for 30 min. Beads were then washed three times in 200 μl Wash buffer. For the second reaction, Template 2 (pG5CG-D2 (Pardee et al., 2003)), wild type yeast nuclear extract (~ 1.5 mg), and ^32^P-labeled NTPs (250 μM of ATP, 250 μM of CTP, 10 μM of UTP and 1.2 μCi ^32^P-labeled UTP) in 240 μl 1X transcription buffer were added to the Template 1-bound beads. At each indicated time point, 60 μl of the mix was collected and the reaction stopped with 200 μl RNAse T1 mixture (1,000 unit, Thermo Scientific). Transcripts were recovered and analyzed by standard gel electrophoresis, autoradiography, and phosphorimager.

## Acknowledgements

We thank Joe Reese for Taf1 antibodies and strains; Kevin Struhl, Fred Winston, Steve Hahn, Frank Holstege, and members of the Buratowski and Marto labs for helpful discussions. This work was supported by DFCI/Blais Proteomics Center support to J.A.M. and NIH grant GM46498 to S.B.

